# Pre-cheliceral region patterning in a spider provides new insights into the development and evolution of arthropod neurosecretory centres

**DOI:** 10.1101/2025.11.10.687188

**Authors:** Amber Harper, Lauren Sumner-Rooney, Ralf Janssen, Alistair P. McGregor

## Abstract

Comparing head development among arthropods has helped identify ancestral aspects of brain patterning and structure in these animals and more broadly. Most understanding of arthropod head patterning has been learned from insects and the myriapod *Strigamia maritima*. Chelicerates represent an outgroup to mandibulate arthropods and can provide a valuable perspective to arthropod evolution and development. We assayed the expression of key markers of head patterning and neurosecretory centres in mandibulates in the pre-cheliceral region of embryos of the spider *Parasteatoda tepidariorum*. We found that, like mandibulates, this spider likely has a *pars intercerebralis*, marked by *six3.2* and *visual system homeobox/chx*. We also found some evidence for another neurosecretory centre, the *pars lateralis,* marked by *six3.2* and *fasciclin 2*. Furthermore, we identified anterior-medial cells in the spider pre-cheliceral region that express *six3.2*, *foxQ2*, and *collier1*, suggesting they may be pioneer neurons. However, these spider cells do not appear to be equivalent to the central pioneer neuronal cells identified in *S. maritima* because they lack expression of other key markers. Taken together, our study of spider pre-cheliceral region patterning adds a new chelicerate perspective to understanding the development and evolution of the arthropod head.

## Introduction

The brain of arthropods is composed of three conserved developmental units, the protocerebral, deutocerebral and tritocerebral ganglia, that together make up the tripartite brain (1, 2) (Fig. 1). The deuterocerebrum is associated with the antennal segment of mandibulates (and the cheliceral segment of chelicerates), and the tritocerebrum is associated with the second antenna-bearing segment of crustaceans (which is appendage-less in insects and myriapods) and the pedipalpal segment of chelicerates (Fig. 1). The anterior-most region of the arthropod head, harbouring the protocerebrum, is either associated with a non-segmental region, or one or two cryptic segments (discussed in e.g. 3, 4-10).

**Figure 1.**
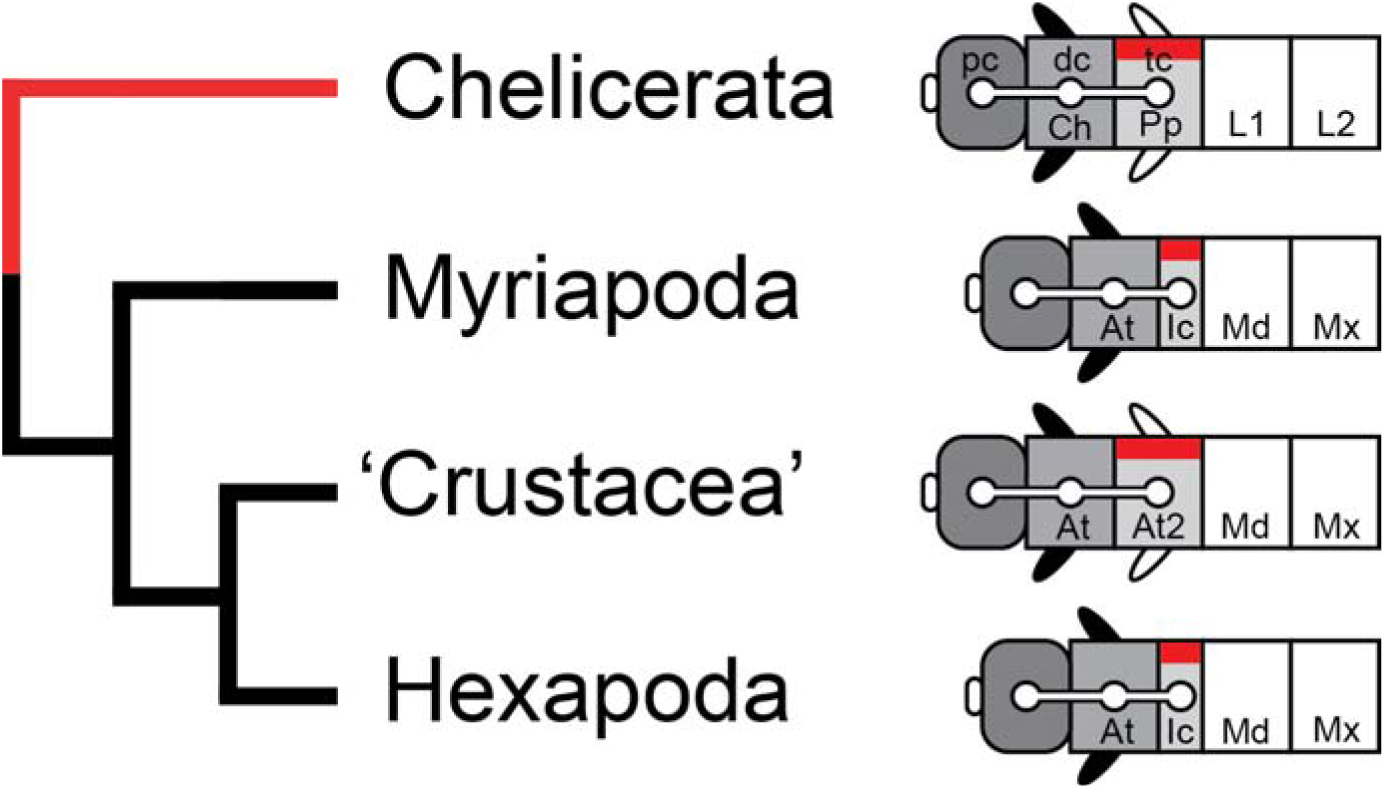
Head structure in arthropods. Schematics of the anterior body region of chelicerates and mandibulates (black branches) with the anterior to the left. Different shades of grey indicate their shared ancestry. The Hox gene *labial* (red) marks the tritocerebral segment. Supraesophageal ganglion: protocerebrum (pc), deutocerebrum (dc), and tritocerebrum (tc). The labrum is shown as an anterior medial structure of the pre-antennal/pre-cheliceral region. At: antennal segment; At2: second antennal segment; Ch: cheliceral segment; Ic: intercalary segment; Md: mandibular segment; Mx: maxillary segment; Pp: pedipalpal segment; L1, L2: walking leg segments L1, L2. Head structure drawings adapted from (11).

In the protocerebrum of *Drosophila melanogaster*, adjacent bilateral placodes expressing *six3* give rise to two neurosecretory centres, the *pars intercerebralis* (PI) and *pars lateralis* (PL), which form parts of the insect neuroendocrine system (10, 12–15).

Neurosecretory cells (NSCs) of the PI and PL project their axons towards endocrine glands, the *corpora cardiaca* and *corpora allata*. In *D. melanogaster*, the *corpora cardiaca*, *corpora allata*, and the prothoracic gland form a single complex, the ring gland, which surrounds the anterior tip of the aorta. The insect PI-PL/ring gland complex is required for several physiological functions critical in development, energy metabolism, growth, water regulation, and reproduction (16, 17).

In insects the PI placode and its derivative neurons express *six3* and *vsx1/chx1* and can be followed until the late pupal stage (12, 13). The PL placode and NSCs of the PL are marked by *six3* and the adhesion protein Fasciclin 2 (Fas2) (12, 13). In the red flour beetle *Tribolium castaneum*, *vsx/chx* and *fas2* expression in the presumptive PI and PL primordia were lost after *six3* knockdown, which suggests Six3 functions relatively early in neuroendocrine development (13). *tailless* (*tll*) also functions upstream in the regulation of neuroendocrine system development in *D. melanogaster* (12). In this insect, *tll* expression encompasses both the PI and PL primordia and loss of *tll* removes most of the protocerebrum including the PI (12). The anterior expression of *foxQ2* (*fd102C*) also appears to locate to the PI and PL in *D. melanogaster* (18). In *T. castaneum*, *chx* initially largely overlaps with *foxQ2*, but they later resolve to mainly non-overlapping patterns (19). Knockdown of *T. castaneum foxQ2* results in the loss of *chx* expression in the early elongating germ band and reduces expression in PI primordia at later stages (19).

Beyond insects, additional key insights to the development and evolution of the anterior head regions and neurosecretory centres of mandibulates have been provided by analysis of the myriapod *Strigamia maritima*. In this centipede, *foxQ2* expression is entirely nested in the *six3*-expressing domain (20). The centipede PI develops within this domain from *vsx/chx*-expressing placodal-like invagination sites (20) like in *D. melanogaster*. However, as the expression of the *S. maritima fas2* ortholog has not been assayed, it remains unclear whether this centipede has a PL as defined by *six3* and *fas2* expression, as in *D. melanogaster* (12, 21).

In addition to the PI, central pioneer neuronal cells (CPNCs) have also been described in *S. maritima*, which have not yet been found in any other arthropod to our knowledge (20). These centipede CPNCs form a dense cluster and, once internalised, project axonal tracts that pioneer the primary axonal scaffold of at least the anterior CNS (20). The *S. maritima* CPNCs arise in the domain of overlapping *six3* and *foxQ2* expression and are characterised by the expression of *collier* (*col*) and *prohormone convertase 2* (*PC2*; also *PHC2*) as well as *nkx2.1*/*scarecrow* (*scro*), *homeobrain* (*hbn*), and *retina and anterior neural fold homeobox* (*rax*), which distinguish them from the PI (20). *col* marks post-mitotic cells committed to a neural fate, while *PC2* encodes an enzyme specifically restricted to endocrine and neuroendocrine cells (22) thus, the *S. maritima* CPNCs are thought to form an active early neurosecretory centre (20). Some centipede CPNCs express the hypothalamic neuron markers *ventral anterior homeobox 1* (*vax1*) and *orthopedia* (*otp*), as well as the *oxytocin/vasopressin* ortholog *inotocin-like* (formerly *vtn*) (20). Interestingly, *vax* genes are absent from insect genomes (22) and the insect *oxytocin/vasopressin* ortholog *inotocin* has been lost from the *D. melanogaster* genome (23). Therefore, equivalent cells to centipede CNPCs have not been observed in insects and even some of the markers of these cells have been lost in these animals.

Better understanding head and brain patterning and evolution in arthropods requires analysis of additional species to compare to existing findings in insects and the myriapod *S. maritima*. The chelicerates represent the sister clade to mandibulate arthropods (Fig. 1) and include spiders, scorpions, harvestmen, mites, ticks, and horseshoe crabs. Spiders have proven to be valuable comparative models to understand many aspects of development and evolution in other arthropods (24–26).

In spiders the protocerebrum develops from the pre-cheliceral region (Fig. 1) and forms the optic ganglia, mushroom bodies, and arcuate body (27). Neurogenesis begins with the appearance of point-like depressions of invaginating neural precursor groups of five to nine cells, which directly differentiate into neurons, in the ventral neuroectoderm and pre-cheliceral lobes (28). Single cell RNA-seq analysis in the spider *Parasteatoda tepidariorum* suggested a potential PI cell population based on co-expression of *six3.2* and *vsx* in a cell cluster and the potential overlapping spatial expression of these genes (29). Furthermore, in the central American wandering spider *Cupiennius salei*, postmitotic cells committed to neural fate were previously identified in the medial pre-cheliceral lobes that give rise to the central protocerebrum and could be similar to the centipede CPNCs (20, 30). Therefore, we aimed to verify the presence of a PI in *P. tepidariorum* and investigate whether this spider has a PL and a cell population like the centipede CPNCs by analysing the expression of known marker genes for these developing neurosecretory centres.

We found further evidence for the presence of a PI and indication of a PL in the pre-cheliceral region of this spider. We also identified an anterior-medial cell population in the spider pre-cheliceral region that is marked by *six3.2*, *foxQ2* and *col1*, but these cells did not express any other markers of the centipede CPNCs.

## Materials and Methods

### Spider culture, embryo staging, and fixation

The *P. tepidariorum* culture was maintained at 25°C in a 12h light/12h dark cycle. Embryos were staged according to Mittmann and Wolff (31). Embryos were dechorionated with bleach (sodium hypochlorite, 5% active chlorine, Arcos) and tap water (1:1). Dechorionated embryos were fixed overnight at room temperature (RT) in a two-phase solution of 100% heptane and 37% formaldehyde in 2xPEM (PIPES, EGTA, and MgSO_4_) (2:1:1). After fixation, embryos were washed with ice-cold 100% methanol, rocked for ≥30 minutes at RT and stored at −20°C. The vitelline membranes of fixed embryos were removed with Dumont 5 forceps in 100% methanol.

### Millipede embryo collection, staging and fixation

*G. marginata* embryos were collected, staged and prepared as described previously (32).

### RNA extraction and cDNA synthesis

*P. tepidariorum* embryos of stages 5-14 were pooled and total RNA was extracted using QIAzol Lysis Reagent (79306, Qiagen), following the manufacturer’s guidelines, and stored at −80°C. Total RNA was used for cDNA synthesis using the QuantiTect Reverse Transcription Kit (205311, Qiagen), according to the manufacturer’s guidelines.

### Primer design and PCR

Primer3web (http://primer3.ut.ee; default settings) was used to design gene-specific primers covering part of the coding region of *col1*, *fas2A*, *foxQ2*, *hbn*, *nkx2.1/scro*, *raxA*, *six3.2*, *tll*, *vsx/chx* (Table S1). For in situ hybridisation, a linker sequence was added to the 5’ end of the forward primer (5′-GGCCGCGG-3’) and reverse primer (5′-CCCGGGGC-3’). Note that the protocol of David and Wedlich (33) was used to design primers to generate probes to assay spider expression of *PC2*, *PC1/3*, *raxB* and *otp* as well as *G. marginata fas2* and *PC2*. Gene-specific cDNA fragments for probe synthesis were amplified using the OneTaq 2x Master Mix (M0482, New England Biolabs/NEB). PCR products were run on a 1% agarose gel to verify band size and purified using the NucleoSpin Gel and PCR Clean-up kit (12303368, Macherey-Nagel).

### Probe synthesis and in situ hybridisation

Fragments of the coding region were amplified from cDNA by two rounds of PCR: the first PCR using gene-specific primers (Table S1) and the second PCR using the gene-specific forward primer and 3’UTP-T7 reverse primer (AGGGATCCTAATACGACTCACTATAGGGCCCGGGGC). Digoxigenin (DIG) and Fluorescein (FITC) labelled RNA probes were synthesised using T7 RNA polymerase (10881775001, Roche) and DIG (11277073910, Roche) or FITC (11685619910, Roche) RNA Labelling Mix, respectively, according to the manufacturer’s guidelines.

For colorimetric in situ hybridisation minor modifications were made to the whole-mount in situ hybridisation protocol described in (34) or (35). Post-fixation embryos were gradually moved from 100% methanol to Phosphate Buffered Saline with Tween-20 (0.02%) (PBS-T). The Proteinase K Digestion (steps 4-8) was replaced by two 10-minute washes in PBS-T (34). For steps 18 and 19, embryos were incubated with gentle agitation at RT for 30 minutes and 2 hours, respectively. For *G. marginata* embryos, 7 µl of NBT/BCIP (Roche, 11681451001) was used in 1 ml of staining buffer instead of using BM-Purple.

For double fluorescence ISH (dFISH), the protocol described by Clark and Akam (36) was followed with some modifications. The post-fixation, pre-hybridisation, hybridisation, and probe removal steps follow steps 1-18 in the protocol described by Prpic *et al.* (34) subject to the modifications stated above. Embryos were incubated with 1:2000 Anti-DIG-alkaline phosphatase (AP) (11093274910, Roche) and 1:2000 Anti-FITC-horse-radish peroxidase (POD) (11426346910, Roche) for 2 hours at RT and washed at least four times for 15 minutes with PBS-T. Embryos were incubated with TSA Plus Biotin (NEL749A001KT, Perkin Elmer) for 5-10 minutes, followed by a 90-minute incubation in 1:500 Alexa Fluor 647 Streptavidin conjugate (S32357, ThermoFisher Scientific]) to visualise the POD signal. The AP signal was visualised by a Fast Red reaction (4210, Kem En Tec Diagnostics).

### Imaging and image processing

Embryos were counterstained with DAPI (10236276001, Roche) (1:2000) or SYBR-green (1:10000) in PBS-T, rocked for 20□minutes at RT and stored in PBS-T at 4□°C. Some of the stained embryos were flat-mounted in PBS-T on poly-L-lysine-coated coverslips and mounted in 80% glycerol. Embryos were imaged using a Zeiss AxioZoom V16 or Zeiss LSM800 Confocal with Airyscan. Images were processed using Image J (FIJI) and Adobe Photoshop.

## Results and Discussion

To better understand the specification of neuroendocrine cells during chelicerate development and how this compares to insects and the centipede *S. maritima*, we examined the relative expression of key marker genes during head development in embryos of the spider *P. tepidariorum*.

### *tll* marks the developing protocerebrum

*P. tepidariorum* has a single *tll* ortholog [g18090] (37). Although expression of this gene was previously characterised from embryonic stages 8 to 12 (38), to further characterise the molecular signature of the potential spider PI and PL, we re-examined the expression of *P. tepidariorum tll* and relative to the segmental marker *hedgehog* (*hh*) (39).

We found that *tll* is expressed from stage 6, earlier than previously reported (38), in a stripe at the anterior rim of the germband (Fig. 2A,B), and then in the head lobes (Fig. 2C,D). The early stripe of expression lies anterior to the splitting pre-cheliceral/cheliceral/pedipalpal *hh1* stripe (39) and thus marks the pre-cheliceral region (Fig. 2F). Similarly, *tll* anterior expression in *D. melanogaster* embryos marks the future head region (40). Unlike in *D. melanogaster* (40) and *T. castaneum* (41); however, *tll* is not expressed in the posterior of the spider germ band (Fig. 2A). Subsequently, *P. tepidariorum tll* expression marks the neurogenic regions of the pre-cheliceral lobes laterally to the labrum and stomodeum (Fig. 2D,E,G). This suggests *tll* is generally required for protocerebrum development in arthropods and could encompass spider PI and PL cells, like in *D. melanogaster* and *T. castaneum* (12, 13, 41).

**Figure 2.**
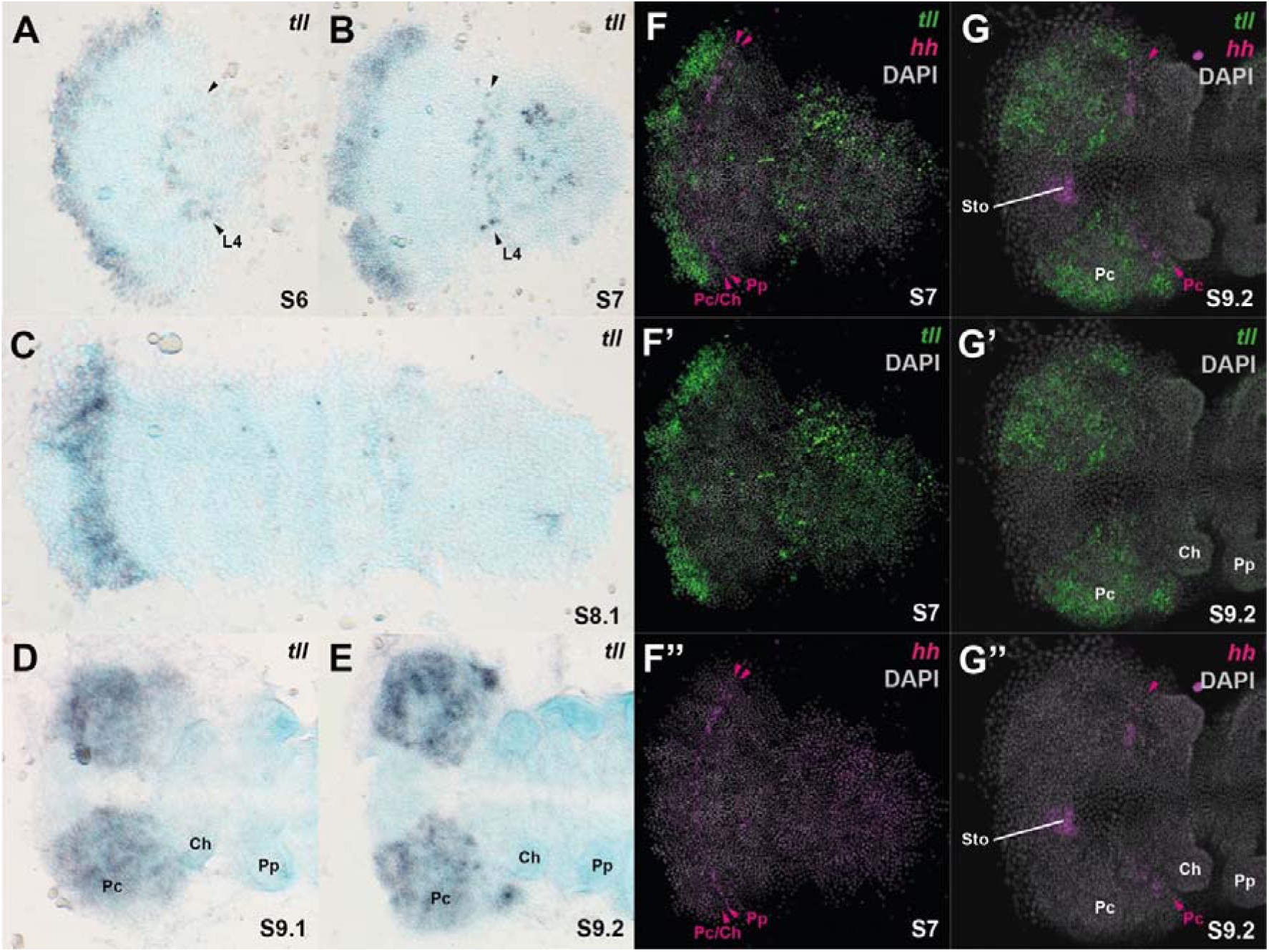
*tll* is expressed in the developing head region of *P. tepidariorum* embryos. **A-E** Embryos stained for *tll* transcripts at stages 6 (**A**), 7 (**B**), 8.1 (**C**), 9.1 (**D**), and 9.2 (**E**). (**F-G’’**) Embryos stained for *tll* (green) and *hh* (magenta) transcripts and nuclei/DNA (DAPI: grey) at stages 7 **(F-F’’)** and 9.2 **(G-G’’)**. **A** *tll* expression was first seen at stage 6 at the anterior rim of the germ band in a broad stripe and in a transient mesodermal stripe (black arrowheads) potentially marking the fourth walking leg segment (L4). **B** Expression at the anterior rim then expands at stage 7. **C** *tll* expression begins to clear from the anterior medial region at stage 8.1. **D,E** Expression splits into two domains and marks the lateral neurogenic regions of the pre-cheliceral region at stages 9.1 and 9.2. **F-F’’** At stage 7, *tll* is expressed anterior to the splitting *hh1* stripes (magenta arrowhead). **G-G’’** At stage 9.2, *tll* is expressed directly adjacent to the pre-cheliceral *hh1* stripe but is not co-expressed with *hh1* in the stomodeum. Ch: chelicerae; Pp: pedipalps; Pc: pre-cheliceral; Sto: stomodeum. Embryos were flat mounted with the anterior to the left.

### Expression of *vsx/chx* and *six3.2* likely demarcates the spider *pars intercerebralis*

To test the prediction that spiders likely have a PI, based on the co-expression of *P. tepidariorum vsx/chx* ortholog (aug3.g27186) and *six3.2* in a pre-cheliceral cell cluster (29, 42, 43), we assayed expression of these two genes from the start of neurogenesis at stage 10.

The expression of *P. tepidariorum vsx/chx* was previously observed along the ventral midline and in the anterior head region (29, 42). At stage 10.1, *vsx/chx* is expressed in a C-shaped domain marking the outgrowing labrum (Fig. 3A-C). *vsx/chx* expression is nearly entirely nested within the *six3.2*-expressing anterior medial region (Fig. 3D-F), like in *T. castaneum* (13). At stage 10.1, *vsx*/*chx* and *six3.2* are co-expressed in and surrounding invaginating neural precursors (INPs) (Fig. 3D). At stage 10.2, *vsx*/*chx* expression also localises to the medial part of the anterior furrow (Fig. 3E), like in *S. maritima* (20). The combined expression of these two genes in *P. tepidariorum* at a corresponding position to that of insects and the centipede strongly suggests that spiders also have a PI.

**Figure 3.**
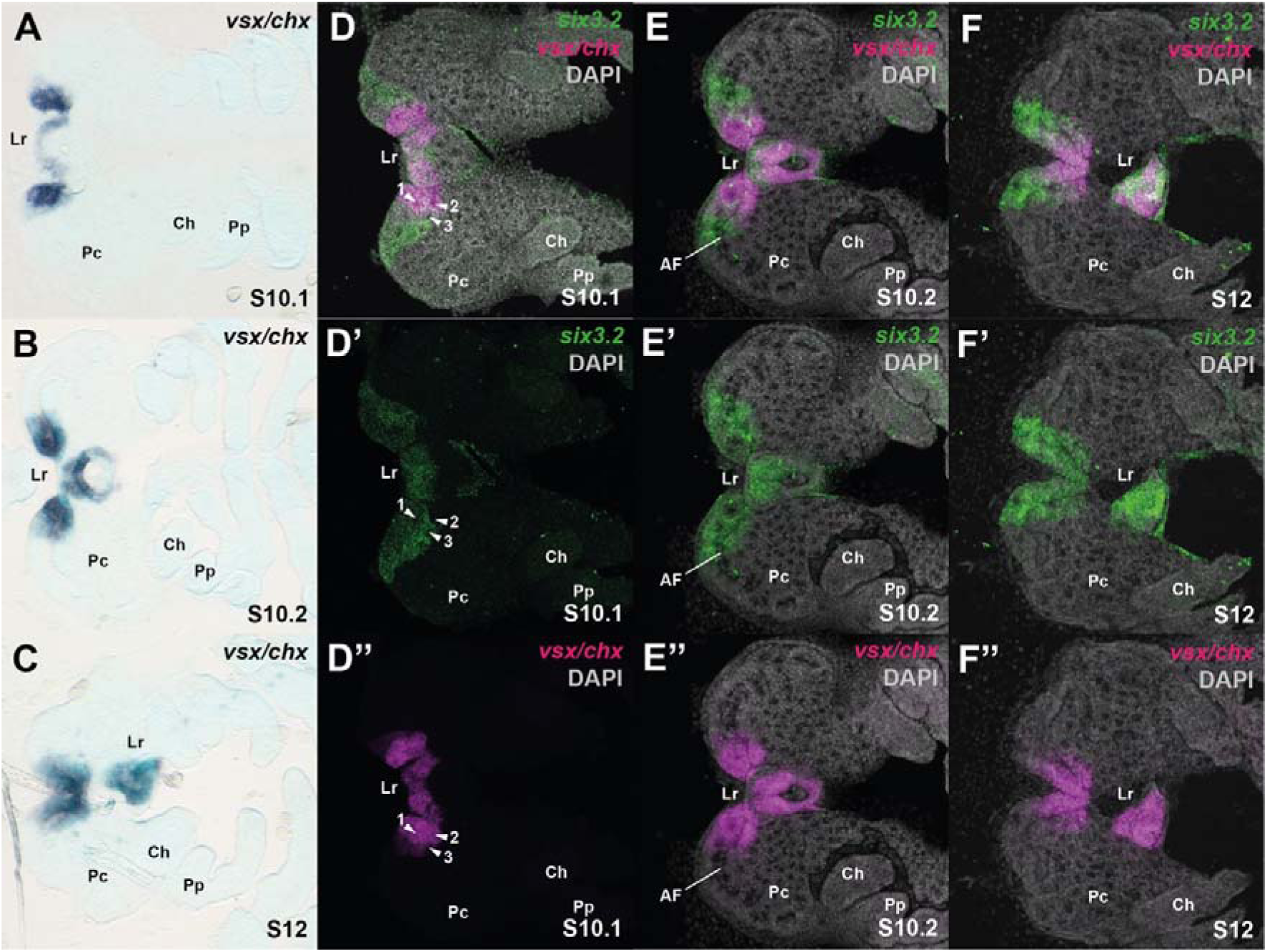
Expression of *vsx/chx* and *six3.2* in the developing head of *P. tepidariorum* embryos. A-C. Embryos stained for *vsx/chx* transcripts at stages 10.1 (**A**), 10.2 (**B**), and 12 (**C**). **D-F’’** Embryos stained for *six3.2* (green) and *vsx/chx* (magenta) transcripts with or without nuclei/DNA (DAPI: grey) at stages 10.1 (**D**), 10.2 (**E**), and 12 (**F**). **D-D’’** Three INPs in the anterior medial region express *six3.2* and *vsx/chx* (1), *vsx/chx* (2), and *six3.2* (3). **E-E’’** At stage 10.2, *vsx* expression localised to the medial part of the anterior furrow (AF). Ch: chelicerae; Lr: labrum; LF: lateral furrow: Pp: pedipalps; Pc: pre-cheliceral region. Embryonic heads were flat mounted with the anterior to the left. *foxQ2* expression is associated with the PI in several panarthropod species (18–20, 44, 45). *P. tepidariorum* has one *foxQ2* ortholog [g224] (46, 47), which is expressed at the anterior rim of the germband at stage 7 and subsequently in several domains in the neurogenic ectoderm, and appears to function downstream of the *six3* genes in this spider (46). We examined the relative expression of *foxQ2* and *vsx/chx* from the onset of *vsx/chx* expression in the putative spider PI. We found that *foxQ2* and *vsx*/*chx* are expressed in abutting, mutually exclusive domains in the developing brain and labrum, although we cannot completely exclude that there is some overlap (Fig. 4). This suggests that while *foxQ2* is expressed, likely downstream of *six3*, early in the presumptive anterior pre-cheliceral region in this spider, expression of this gene is not later associated with the putative spider PI cells as defined by overlapping *six3* and *vsx/chx* expression (Fig. 3). However, the largely non-overlapping expression of *foxQ2* and *vsx/chx* as these patterns resolve during brain development is similar between *P. tepidariorum* and *T. castaneum* (19).

**Figure 4.**
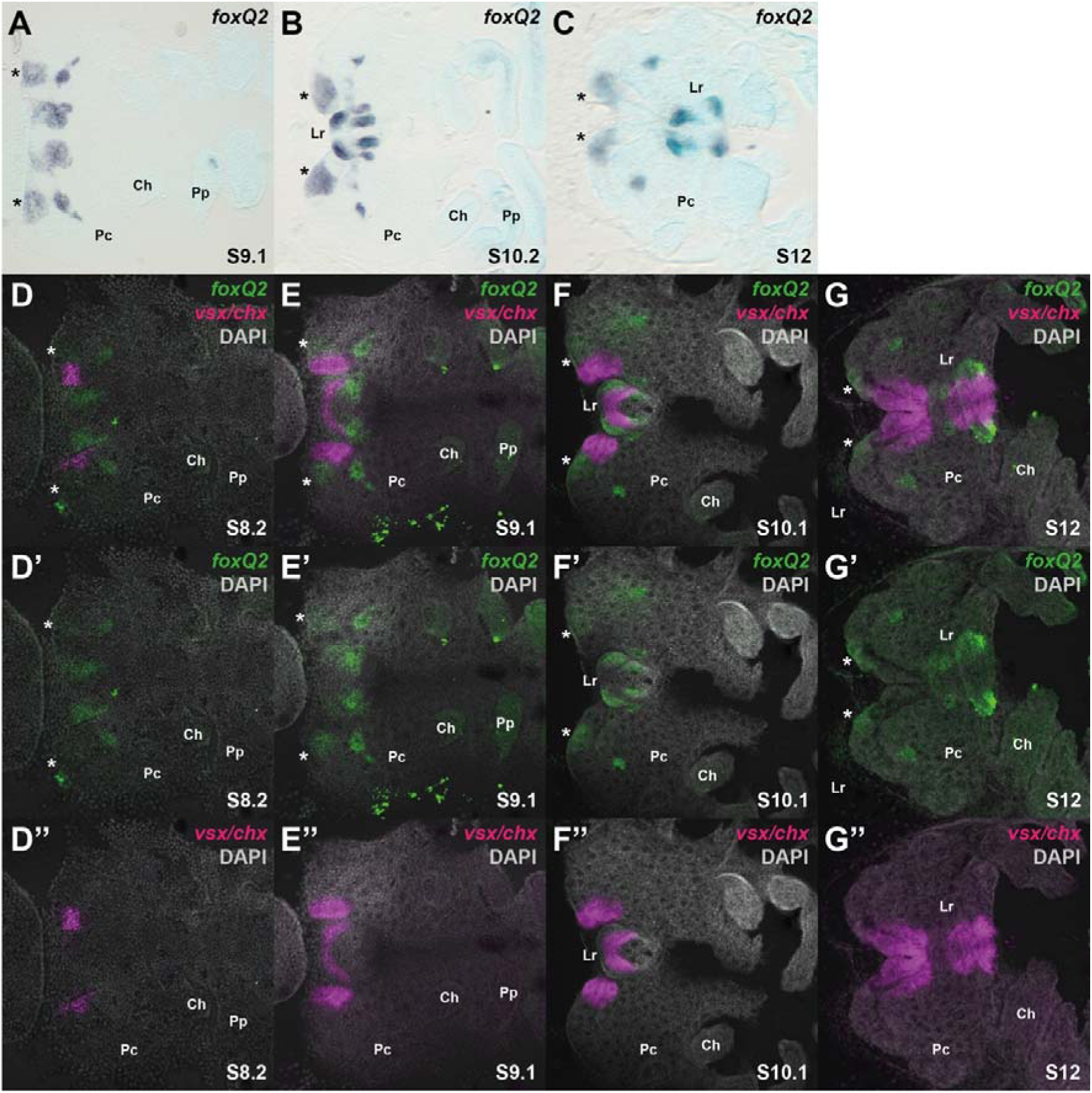
Expression of *vsx/chx* and *foxQ2* in the developing head of *P. tepidariorum* embryos. A-C. Embryos stained for *foxQ2* transcripts at stages 9.1 **(A**), 10.1 (**B**), and 12 (**C**). **D-G’’** Embryos stained for *foxQ2* (green) and *vsx/chx* (magenta) transcripts and nuclei/DNA (DAPI: grey) at stages 8.2 (**D**), 9.1 (**E**), 10.1 (**F**), and 12 (**G**). Ch: chelicerae; Lr: labrum; Pp: pedipalps; Pc: pre-cheliceral. Corresponding expression domains in all panels are indicated by an asterisk. Embryos were flat mounted with the anterior to the left.

### Expression of *fas2A* indicates spiders may have a *pars lateralis*

In insects the PL is marked by Fas2 expression exclusive of *vsx/chx* (12, 13) (Fig. 2). *P. tepidariorum* has two *fas2* orthologs, *fas2A* [g31369] and *fas2B* [g18551], and their expression was previously characterised from embryonic stages 8 to 12 (38). *fas2A* has a complex and dynamic expression pattern including in proneural clusters in the head, while *fas2B* has ubiquitous and uniform expression throughout the embryo (38). To explore whether spiders have a PL, we re-examined the expression of *P. tepidariorum fas2A*.

We observed *fas2A* expression from stage 9.2 in the neurogenic ectoderm of the pre-cheliceral lobes and segmentally repeated domains along the body (Fig. 5), earlier than previously reported (38). From the onset of expression, the anterior-most medial domain overlapped with *six3.2* (Fig. 5E), as in *T. castaneum* (13). At stage 10.1, this spot of *fas2A* expression was observed during the formation of an INP (Fig. 5F). Consistent with *D. melanogaster* (12) and *T. castaneum fas2* (13), expression of *P. tepidariorum fas2A* does not overlap with *vsx*/*chx* (Fig. 5G,H). This suggests the PI expressing *vsx/chx* in *P. tepidariorum*, is distinct from cells expressing *fas2A* and that spiders may have a PL.

**Figure 5.**
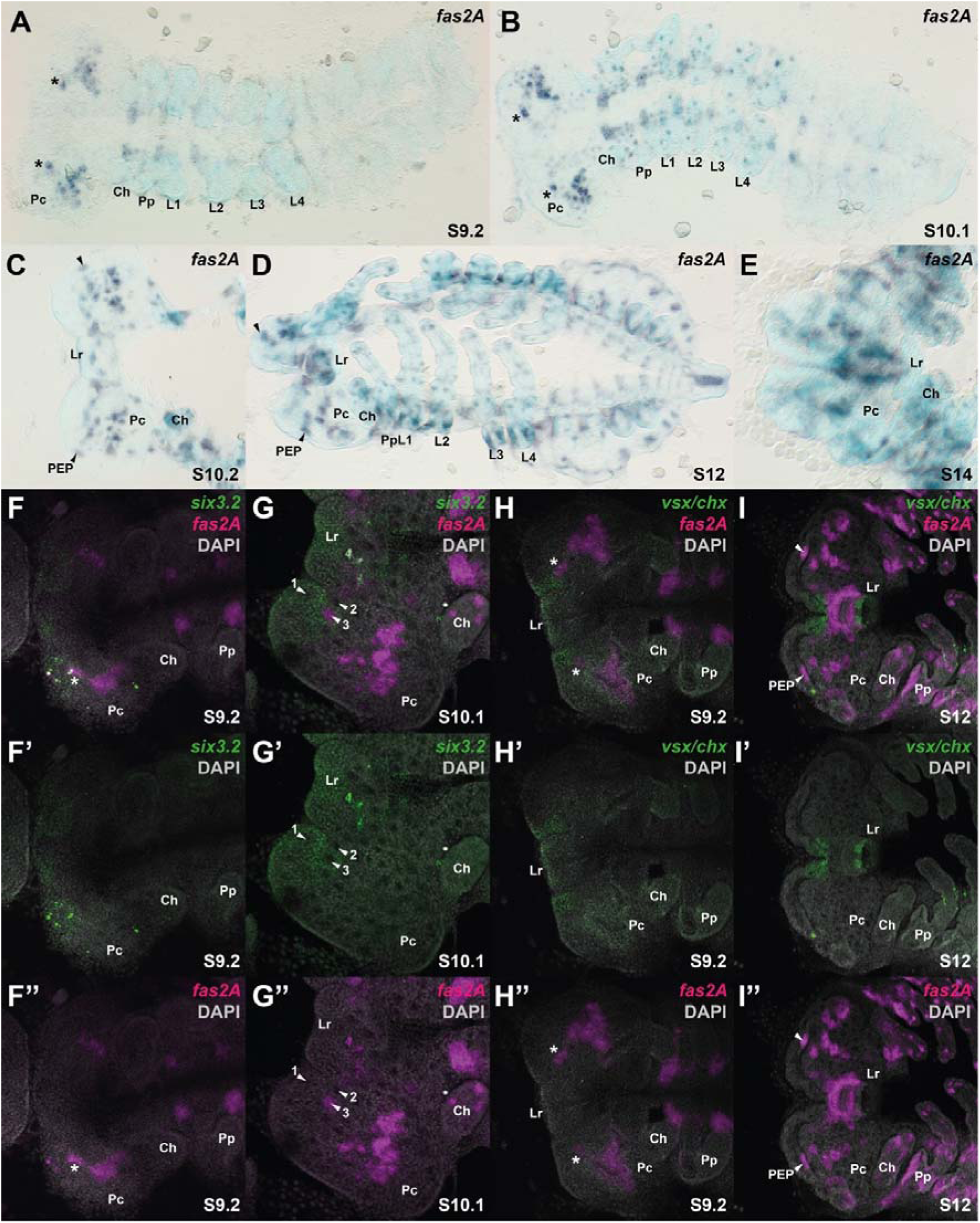
Expression of *fas2A* in *P. tepidariorum* embryos. **A-E** Embryos stained for *fas2A* transcripts at stages 9.2 (**A**), 10.1 (**B**), 10.2 (**C**), 12 (**D**), and 14 (**E**). **F-G’’** Embryos stained for *six3.2* (green) and *fas2A* (magenta) transcripts and nuclei/DNA (DAPI: grey) at stages 9.2 (**F**) and 10.1 (**G**). **G** Three point-like depressions in the anterior medial region express *six3.2* and *vsx/chx* (1, see Figure 4D), *vsx/chx* (2, see Figure 4D), and *six3.2* and *fas2A* (3). **H-I’’** Embryos stained for *vsx/chx* (green) and *fas2A* (magenta) transcripts and nuclei/DNA (DAPI: grey) at stages 9.2 (**H**) and 12 (**I**). Ch: chelicerae; Lr: labrum; Pp: pedipalps; Pc: pre-cheliceral; PEP: principal eye primordia; L1-4: walking legs 1-4. Corresponding expression domains in all panels are indicated by an asterisk. Embryos were flat mounted with the anterior to the left.

Although *fas2* expression has not been described in the centipede *S. maritima*, we assayed expression of this gene during embryogenesis in another myriapod, the millipede *G. marginata* (Fig. S1). We observed expression in anterior medial cells of the developing head of *G. marginata* embryos suggesting myriapods also have a PL and this could therefore be an ancestral neurosecretory centre in arthropods (Fig. S1).

### Other potential roles of *fas2* in *P. tepidariorum*

We also observed that in stage 10.2 *P. tepidariorum* embryos, *fas2A* is expressed in three spots in the dorsal part of the stomodeum (Fig. 5C). In spiders, INPs arranged in a crescent-shaped array in the dorsal part of the stomodeum were suggested to give rise to the stomatogastric nervous system (SNS) (27) like in *D. melanogaster*, where the SNS primordium forms from three invaginating placodes in the roof of the foregut (48). Therefore, we suggest that *fas2A* contributes to the formation of the SNS in the spider. Furthermore, at stage 10.2 there is an additional *fas2A* domain at the rim of the head lobes, adjacent to the anterior furrow (Fig. 5C,D). This *fas2A* expression domain remains at the leading edge of the non-neurogenic ectoderm and could correspond to the principal eye primordia (Fig. 5D).

### Spiders have a *col-*expressing anterior neuronal cell population, but do not appear to possess a cell population equivalent to centipede CPNCs

Centipede CPNCs are characterised by the expression of *col* and *PC2*/*PHC2* nested within a *six3* and *foxQ2* expression domain, and they express *otp* in a subset of cells as well as *nkx2.1*/*scro*, *hbn*, and *rax* in cells around the neural cells (20). To determine whether spiders have a similar cell population, we first examined whether spiders have a neurogenic *six3* and *foxQ2* expression domain. We found that only the anterior-most domain of *foxQ2* expression is nested within the *six3.2* domain (Fig. 6A-D). At stage 10.1, several point-like depressions of internalised neural precursor cells are observed where the expression of *six3.2* and *foxQ2* overlap (Fig. 6C).

**Figure 6.**
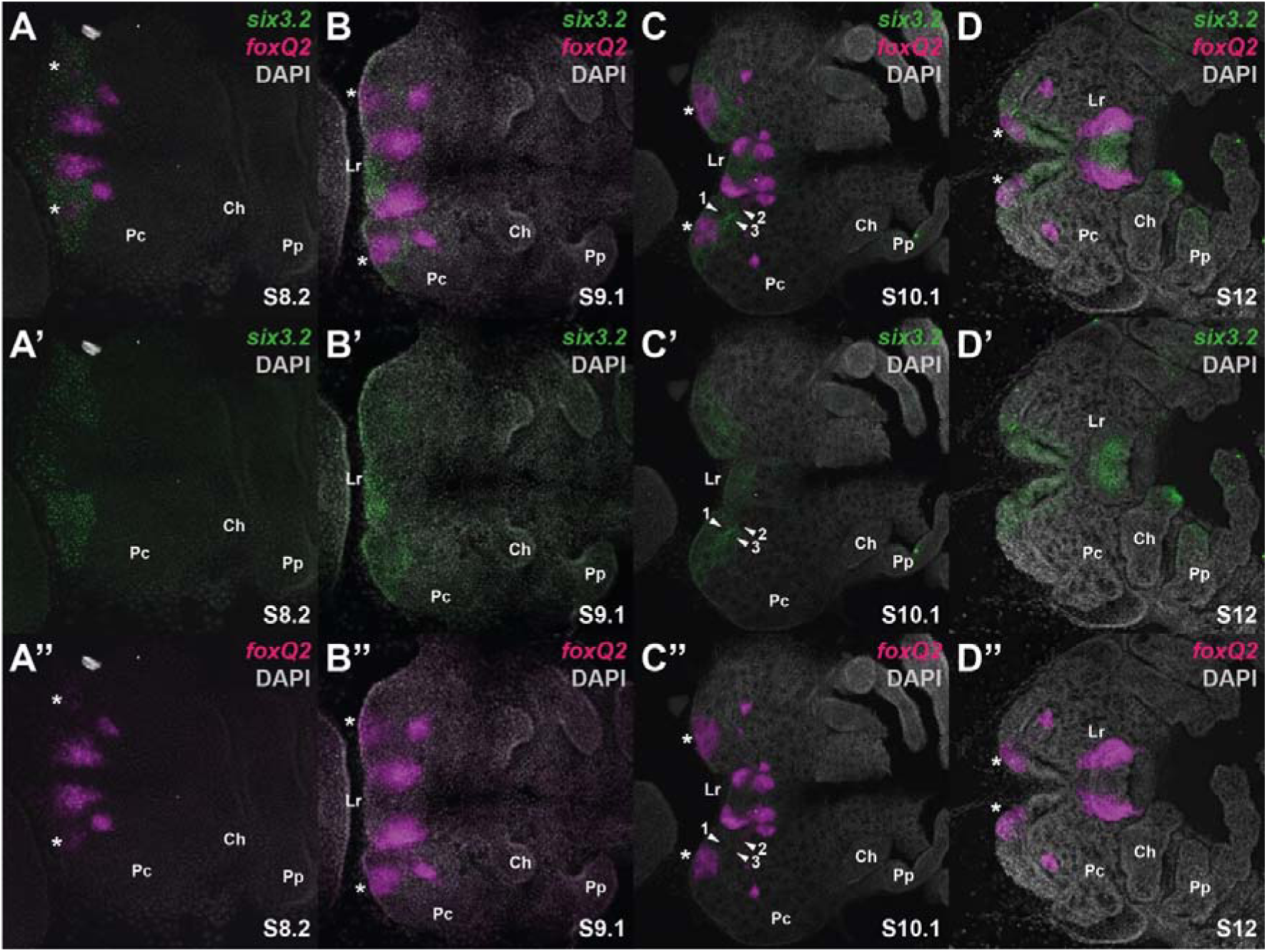
Expression of *six3.2* and *foxQ2* in the developing head of *P. tepidariorum* embryos. **A-D’’** Embryos stained for *six3.2* (green) and *foxQ2* (magenta) transcripts and nuclei/DNA (DAPI: grey) at stages 8.2 (**A**), 9.2 (**B**), 10.1 (**C**), and 12 (**D**). **C** Three point-like depressions in the anterior medial region that express *six3.2* and *vsx/chx* (1, see Figure 3D-D’’), *vsx/chx* alone (2, see Figure 3D-D’’), and *six3.2* and *fas2A* (3, see Figure 5G-G’’) are all free of *foxQ2* expression. Other abbreviations: Ch: chelicerae; Lr: labrum; Pp: pedipalps; Pc: pre-cheliceral. Embryos were flat mounted with the anterior to the left. Corresponding expression domains in all panels are indicated by an asterisk.

*P. tepidariorum* has two *col* orthologs, and their expression was characterised previously at stage 12. *col1* [g10955] was reported to be expressed in a small number of cells at the anterior rim of the head lobes at stage 12 (49), while *col2* [g31501] was described as expressed at stage 12 in bilateral spots in the L4, O1, and O2 segments (38). We re-examined the expression of *col1*.

*col1* is first expressed in the pre-cheliceral lobes at stage 10.2 (Fig. 7A,B). At stages 11 and 12, medial *col1* expression domains at the anterior rim of the pre-cheliceral lobes overlap with *six3.2* (Fig. 7), and likely the anterior-most expression domain of *foxQ2* (Fig. 5D). This indicates that the spider pre-cheliceral region contains *col1* expressing cells, likely committed to a neural fate, nested within a *six3/foxQ2* expression domain, as observed in centipede CPNCs (20). Interestingly, *col*-expressing anterior-medial cells are also observed in embryos of another centipede, *Lithobius forficatus*, as well as the millipede *G. marginata* and the onychophoran *Euperipatoides kanangrensis*, suggesting this pattern is ancestral in panarthropods (50).

**Figure 7.**
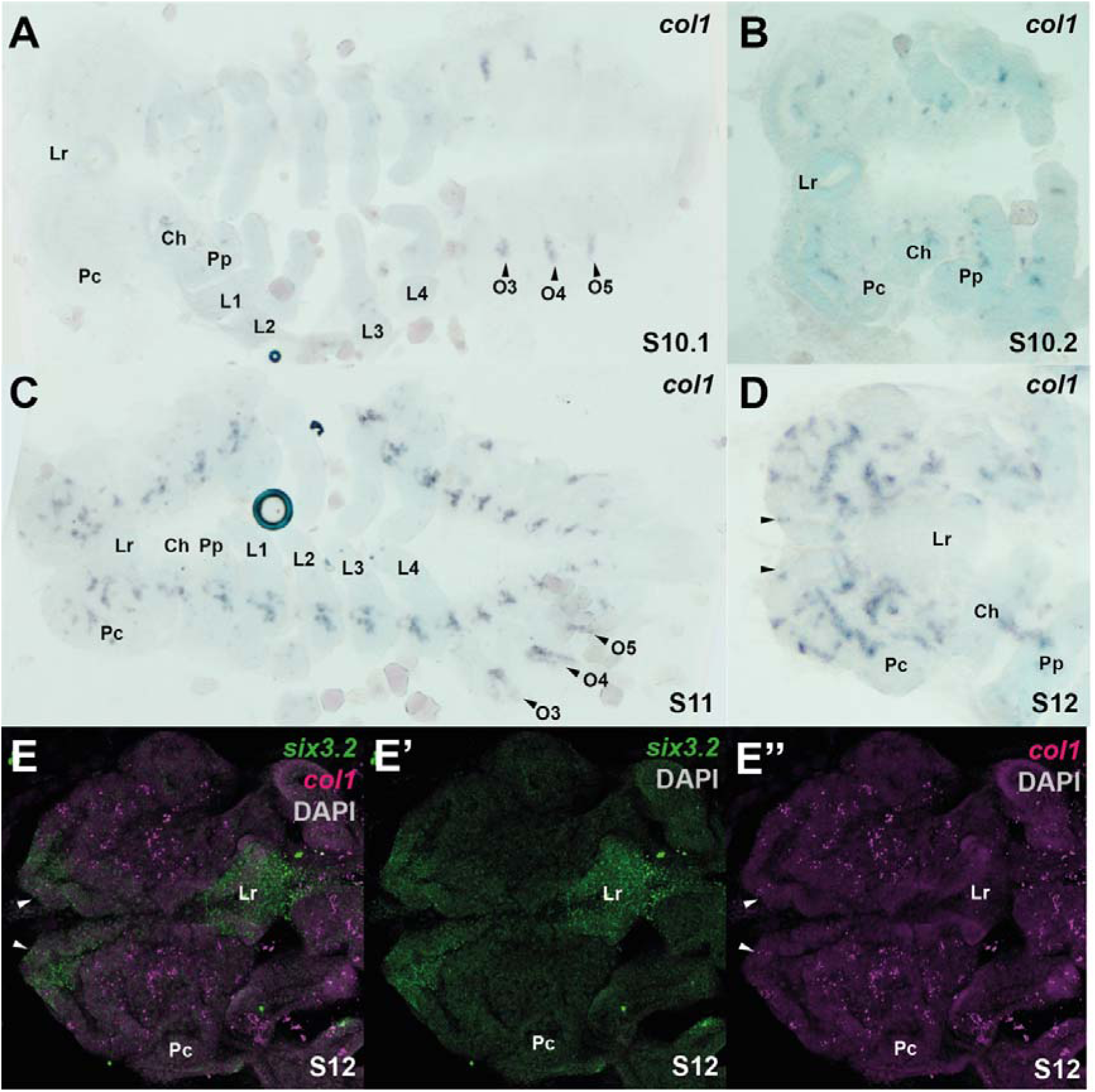
Expression of *col1* in the developing head of *P. tepidariorum* embryos. **A-G** Embryos stained for *col1* transcripts at stages 10.1 (**A**), 10.2 (**B**), 11 (**C**), and 12 (**D**). **A** *col1* expression was first seen at stage 10.1 in the anterior boundary of the third, fourth, and fifth opisthosomal segments (O3, O4 and O5; black arrowheads). **B**. Expression was first observed in the pre-cheliceral region at stage 10.2. By stage 11 and 12 expression was seen in cells at the anterior of the germband (**C,D**)**. E-E’’** Embryos stained for *six3.2* (green) and *col1* (magenta) transcripts and nuclei/DNA (DAPI: grey) at stage 12. At stage 12, arrowheads indicate anterior-medial cells where expression of *col1*, *six3.2* and *foxQ2* overlap (see Figure 6D-D’’). Ch: chelicerae; Lr: labrum; Pp: pedipalps; Pc: pre-cheliceral. Corresponding expression domains in **D** and **E-E’’** are indicated by arrowheads. Embryos were flat mounted with the anterior to the left.

To explore the identity of these spider anterior-medial cells further, we looked at the expression of the *P. tepidariorum* orthologs of *PC2*, *nkx2.1/scro*, *hbn*, *rax and otp*, which also mark the centipede CPNCs.

*P. tepidariorum* has a single *PC2* ortholog [g24083] (51) but we observed that this gene is only expressed in late embryogenesis, from stage 13.1. At this stage *PC2* expression appears in the developing brain (Fig. S2) and subsequently, at stage 13.2, additional dots of expression appear in the appendages and dorsally on either side of the developing dorsal tube (heart) of the spider (Fig. S2).

Likewise, the *T. castaneum PC2* ortholog is first expressed only during late embryogenesis in the developing brain and ventral neuroectoderm, and subsequently in larval nervous system with strong expression in the anterior medial brain (51). To test if early expression of *PC2* associated with CPNCs was more generally found in myriapods, we assayed the expression of this gene in *G. marginata* (Fig. S1). We found that like the insect and spider *PC2* orthologs, *G. marginata PC2* is not expressed during early embryonic stages (Fig. S1).

We then assayed the expression of a related gene, *PC1/3* [XP_015924123.1], in *P. tepidariorum* (51). This gene is first expressed transiently at stage 10.2 at the interface of the opisthosoma of the embryo (proper) and the adjacent so-called extraembryonic tissue (Fig. S2). This expression is possibly associated with the developing heart. Later, at stage 13.2, *de novo* expression appears as domains in each brain hemisphere that expand ventrally and posteriorly forming a bean-shaped pattern (Fig. S2). Therefore expression of *PC1/3* in the developing brain only appears later than the co-expression of *six3.2*/*foxQ2*/*col1* at stage 10.2 in presumptive neural cells in the pre-cheliceral region. Although *PC1/3* has a pre-bilaterian origin, it has been lost in *D. melanogaster* and the only arthropod ortholog that has been investigated is that of *T. castaneum*, which is not expressed in the protocerebrum (51). The single spider ortholog of *nkx2.1/scro* (37, 42, 43) is expressed in the anterior neuroectoderm in a circular domain at stage 8.1 and at stage 10.1 appears to overlap with the posterior part of the *vsx/chx* expression domain that corresponds to the presumptive spider PI (Figs 3,4 and S3). This gene is later expressed around the stomodeum and abutting the anterior and lateral furrows, but no expression was observed in the medial anterior neuroectoderm where *six3.2*, *foxQ2* and *col1* are co-expressed (Fig. S3).

In *D. melanogaster*, *nkx2.1/scro* is expressed in bilateral domains in the developing protocerebrum (52) and in the larval CNS including the PI region (53). In *T. castaneum*, some cells of the developing PI appear to express *nkx2.1/scro* (13), as we report in the spider. In *S. maritima*, *nkx2.1* was described as absent from the PI (20); however, there are lateral dots of expression in this centipede that look similar to expression in *G. marginata* (44, 45), which may correspond to the PI and suggest possible conservation of this *nkx2.1* expression among arthropods.

*P. tepidariorum hbn* [g8762] (37) is first expressed at stage 8.1 in the anterior-lateral neurogenic ectoderm of each pre-cheliceral lobe. From stage 10.2, this gene is expressed within and surrounding the lateral part of the anterior furrow and the labrum (Fig. S4). However, no *hbn* expression is observed in the medial anterior neuroectoderm of *P. tepidariorum* during any stages assayed (Fig. S4).

Spiders have two *rax* orthologs, *raxA* [g8760] and *raxB* [g23262] (37, 43)*. raxA* is expressed from stage 8.1 in a patch in the anterior-lateral neurogenic ectoderm of each pre-cheliceral lobe (Fig. S5). This pattern later subdivides into three domains in and surrounding the lateral part of the anterior furrow, the medial pre-cheliceral lobes, and lateral of the lateral furrow (Fig. S5). *raxB* is only expressed in the developing brain of late-stage embryos (from stage 13.1) (Fig. S6). Therefore neither spider *rax* ortholog overlaps with *six3.2*, *foxQ2* and *col1* at stage 10.2 in the antero-medial region of the pre-cheliceral region, which is consistent with the absence of the expression of this gene in the medial forebrain of insects.

Finally, *P. tepidariorum* has a single *otp* gene [g8767] (37, 43). Despite multiple attempts, we did not detect any embryonic expression of *otp* in this spider. We assume, however, that it is expressed during later embryonic stages, when cuticle formation prevents detection via in situ hybridisation, because we could isolate it from cDNA from embryos including stage 14. Thus, it is unlikely that *otp* is involved in neuroendocrine development in spiders.

Taken together, although we found co-expression of *six3.2*, *foxQ2*, and *col1* in the anterior-medial neuroectoderm of *P. tepidariorum*, expression of these genes did not overlap with *PC1/3*, *PC2*, *nkx2.1/scro*, *hbn*, *raxA*, *raxB* or *otp*, the putative markers of the CPNCs in *S. maritima*. Furthermore *PC2* is only expressed in the developing brain of *G. marginata* during later embryogenesis. These data suggest that, like insects, spiders and millipedes do not possess an active early neurosecretory cell population equivalent to the centipede CPNCs. Indeed, the long-range pioneer neurons originating from the *S. maritima* CPNCs have not been characterised in any other arthropod (20). However, since the centipede CPNCs have a similar transcriptional profile to the apical plate of marine larvae, for example in the annelid *Platynereis dumerilii*, these cells may have a deep evolutionary origin in animals (54–56). This would suggest that the equivalent cell population has been lost in insects and chelicerates. Therefore it would be interesting to investigate the expression of markers of these cells in other groups of chelicerates and myriapods as well as other panarthropods.

In most insects, neuroblasts that pioneer the axon tracts of the anterior CNS are located in the bilateral head neuroectoderm (13, 57). In the grasshopper *Schistocerca gregaria*, two neurons that originate directly from the epithelium within the anterior medial domain of the head pioneer the primary brain commissure (58). The spider anterior-medial *col1*-expressing pre-cheliceral cells identified here may not form an active early neurosecretory centre, but they may correspond to the long-range pioneer neurons of the centipede and/or short-range pioneer neurons of the grasshopper. Consistent with this hypothesis, the later expression of spider *col1* suggests this gene is involved in establishing the anterior commissure (Fig. 7).

## Conclusions

We assayed the expression of markers of neural cell populations identified in insects and a centipede in a chelicerate, the spider *P. tepidariorum*, in order to understand the patterning and evolution of the head and brain across arthropods. Further evidence of a PI in this chelicerate (Fig. 8) suggests this neuroendocrine centre date back to at least the arthropod common ancestor and perhaps even has a panarthropod origin, based on the anterior expression of *six3* and *vsx/chx* in embryos of *E. kanangrensis* (15, 44). The origins of the PL remain more enigmatic. *fas2A* expression in *P. tepidariorum* represents tentative evidence that spiders have a PL like insects (Fig. 8) and our data suggest this could also be the case in the millipede *G. marginata*. However, it will be crucial to identify where *fas2* expression localises during head development in *G. marginata* and other myriapods as well as onychophorans to determine whether a PL is also present in these groups, and thus could also have a panarthropod origin. Finally, we observed a *col-*expressing anterior-medial neuronal cell population embedded in a *six3* and *foxQ2*-expressing domain in *P. tepidariorum* (Fig. 8). This is consistent with the expression of *col1* in other arthropods and the CPNCs in *S. maritima* (20, 50), although these spider cells do not express any other CPNC markers. Thus spiders, like insects, do not appear to have a cell population equivalent to the centipede CPNCs.

**Figure 8.**
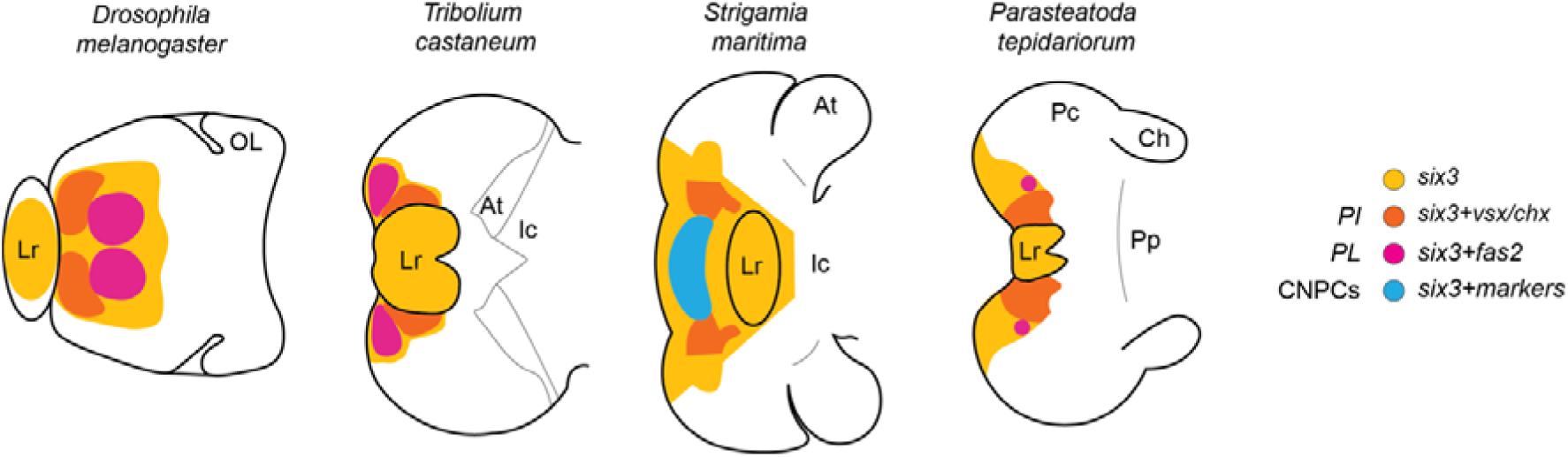
Summary of anterior neurosecretory centres found in different arthropods. Head schematics are orientated with the anterior to the left depicting neuroendocrine cell populations and their molecular signatures defined in *D. melanogaster*, *T. castaneum* and *S. maritima*, and putative homologous clusters in *P. tepidariorum*. *PI*: *pars intercerebralis*. *PL*: *pars lateralis*. CPNCs: central pioneer neuronal cells (marked by *six3* + *PC2*, *nkx2.1/scro*, *hbn*, *rax* and *otp*). OL: optic lobe. At: antennal segment. Ic: intercalary segment. Ch: cheliceral segment. Pp: pedipalpal segment. Lr: labrum. Pc: pre-cheliceral region. Schematics adapted from (13, 15, 20). For the labrum only *six3* expression is indicated. Note that *fas2* expression has not been investigated in *S. maritima*.

## Acknowledgements

We thank Vera Hunnekuhl and Nico Posnien for their comments on the manuscript.

## Authors’ contributions

The project was conceived by AH under the supervision of APM and LSR. Experiments were carried out by AH and RJ. All authors contributed to data analyses and interpretation. The manuscript was drafted by AH and APM and all authors contributed to the final version.

## Data availability

All data generated or analysed during this study are included in this article and its supplementary files.

