## Supplementary Figures 1-6 and Table 1 for "Pre-cheliceral region patterning in a spider provides new insights into the development and evolution of arthropod neurosecretory centres"

**Supplementary Figures and Table**

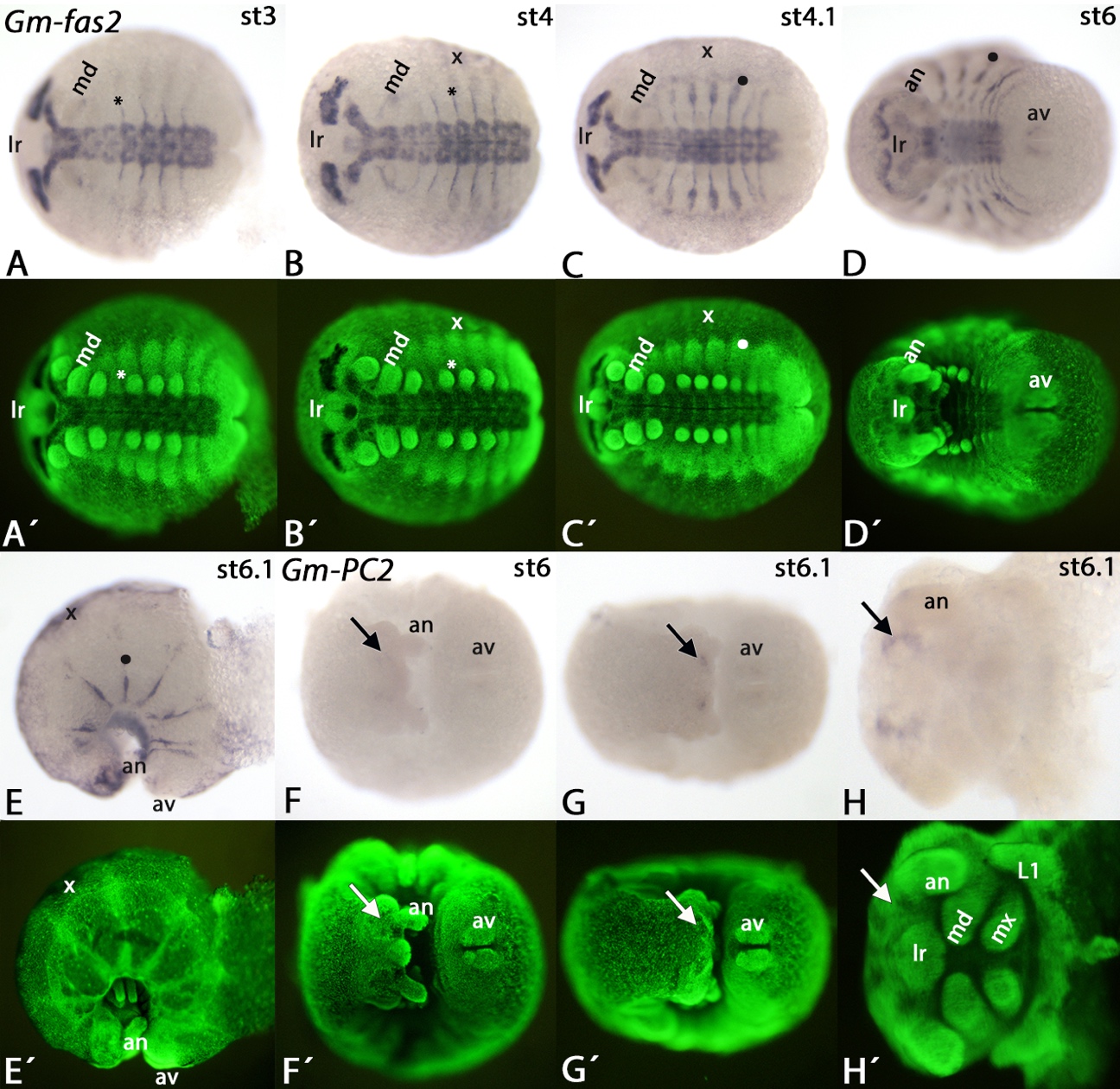

**Figure S1. Expression of *Glomeris marginata fas2* and *PC2*.**

**A**-**E** *fas2* is expressed in the developing ventral nervous system and the developing brain, but is not expressed in the posterior region of the segment addition zone or the appendages (**A**-**D**). Additional expression, however, is seen as small stripes anterior and dorsal to the base of the legs (**A**,**B**; asterisks). Later, these stripes expand towards the dorsal and lie in the middle of the dorsal segmental units (**C**,**D**; filled circles) (cf. Janssen et al. 2004). At late developmental stages, faint expression appears in the anal valves (**D**,**E**), and from approximately stage 4 onwards also in the so-called extra-embryonic tissue (**B**-**E**, marked by “x”). **F**-**H** *PC2* is exclusively expressed the developing brain during late developmental stages when the germ band has already bent inwards (**F**-**H**; arrows). Additional faint expression is inside the anal valves (**F**,**G**). In all panels, embryos are located with anterior to the left and ventral views (except for panel **E** which shows a lateral view). Panel **H** shows magnification of a dissected head (ventral view). Panels with an apostrophe represent SYBR-green staining of the embryos shown in corresponding panels. Abbreviations: an, antenna; av, anal valves; md, mandible; mx, maxilla; L1, first pair of legs; lr, labrum.

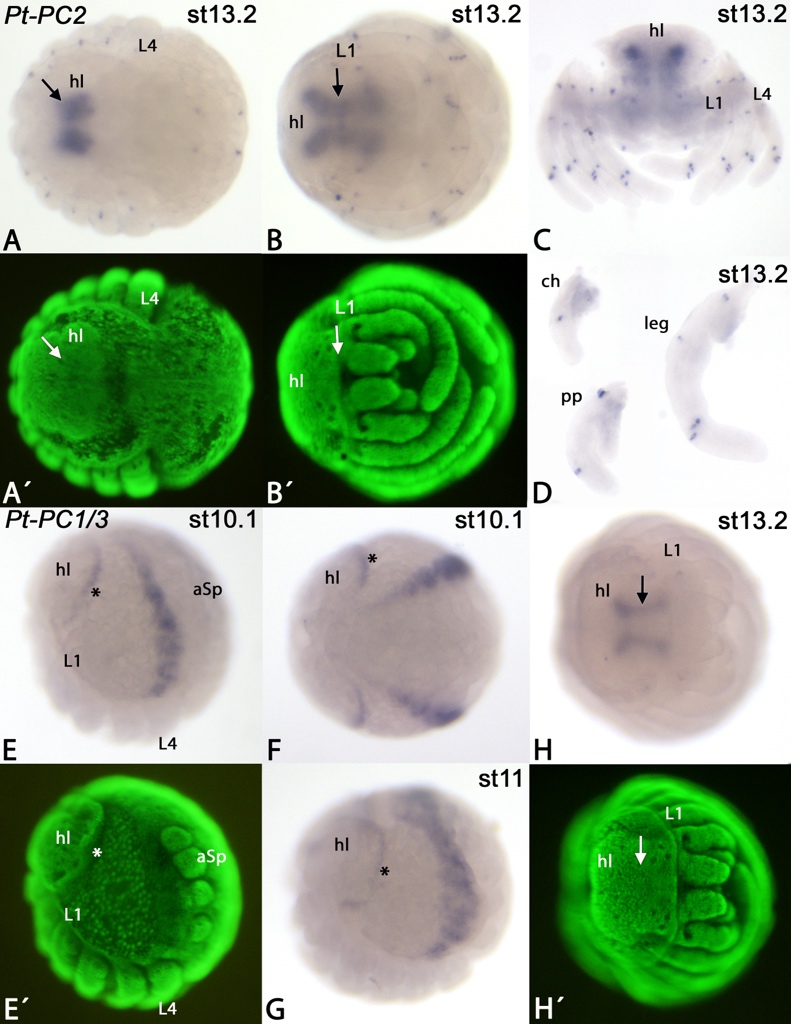

**Figure S2. *Expression of Parasteatoda tepidariorum PC2* and *PC1/3.***

Developmental stages are indicated in the upper right corner. Panels with an apostrophe represent SYBR Green staining of embryos in the corresponding panels. The arrows in panels **A** and **B** point to expression of *PC2* in the developing brain. Note the distinct spots of *PC2* expression that represent single cells of the appendages (**A**-**D**) and flanking the dorsal tube (**A**). Panel **D** shows dissected appendages and the arrow points to expression of *PC1*/3 in the brain. The asterisks in panels **E** and **F** mark *PC1/3* expression anteriorly abutting the head lobes. Abbreviations: aSP, anterior spinneret; ch, chelicera; hl, head lobe; L, leg; pp, pedipalp.

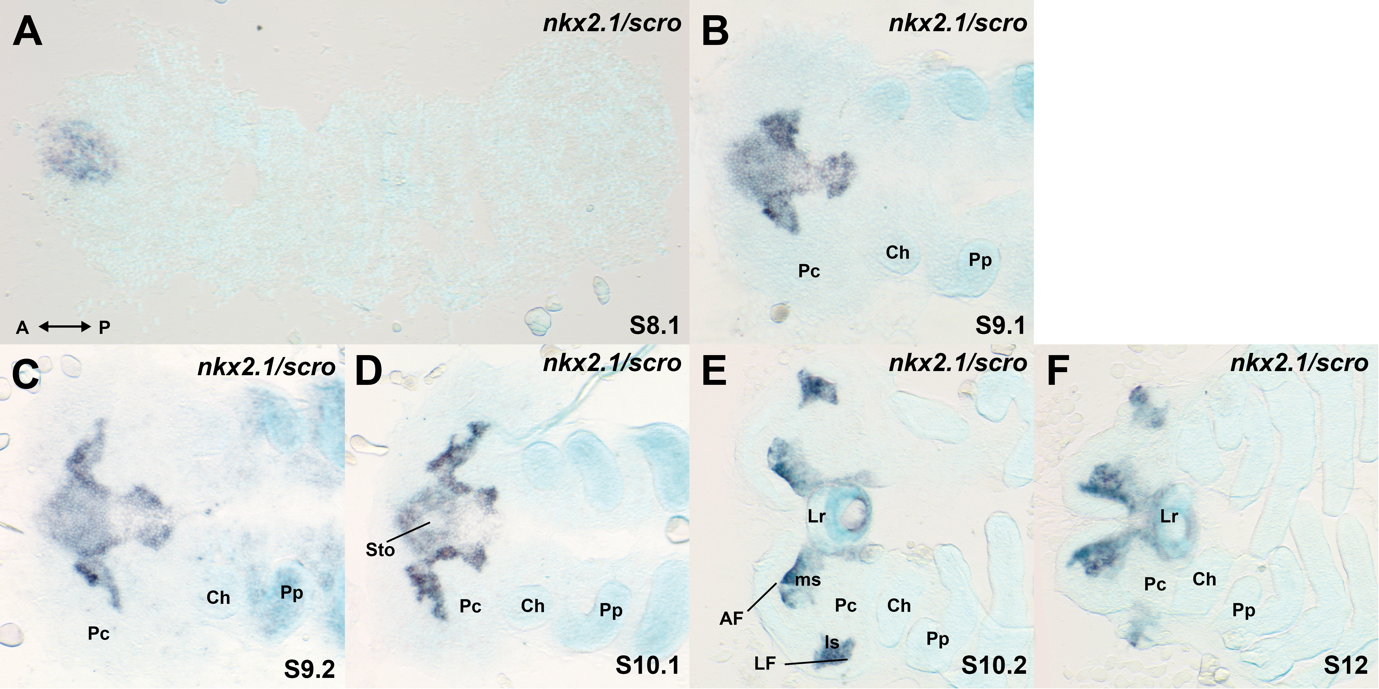

**Figure S3. Expression of *nkx2.1/scro* in the developing head of *P. tepidariorum* embryos.**

**A-F** Embryos stained for *nkx2.1/scro* transcripts at stages 8.1 **(A)**, 9.1 **(B)**, 9.2 **(C)**, 10.1 **(D)**, 10.2 **(E)**, and 12 **(F).** **A** *nkx2.1/scro* expression was first seen at stage 8.1, in a circular domain at the anterior rim of the germ band. **B-D** From stage 9.1, *nkx2.1/scro* expression retracts from the anterior rim and extends from the circular domain into the neurogenic ectoderm. **D** At stage 10.1, *nkx2.1/scro* expression clears at the centre of the circular domain, resulting in expression surrounding the stomodeum (Sto). **E** At stage 10.2, *nkx2.1/scro* is expressed in the medial subdivisions (ms) abutting the anterior furrow (AF) and an additional expression domain arises in the lateral subdivisions (ls) abutting the lateral furrow (LF). **F** At stage 12, the AF and LF were closed by the expansion of the respective subdivisions and the two halves of the brain fuse at the midline. Other abbreviations: Ch: chelicerae; Lr: labrum; Pp: pedipalps; Pc: pre-cheliceral. Embryos were flat mounted with the anterior to the left.

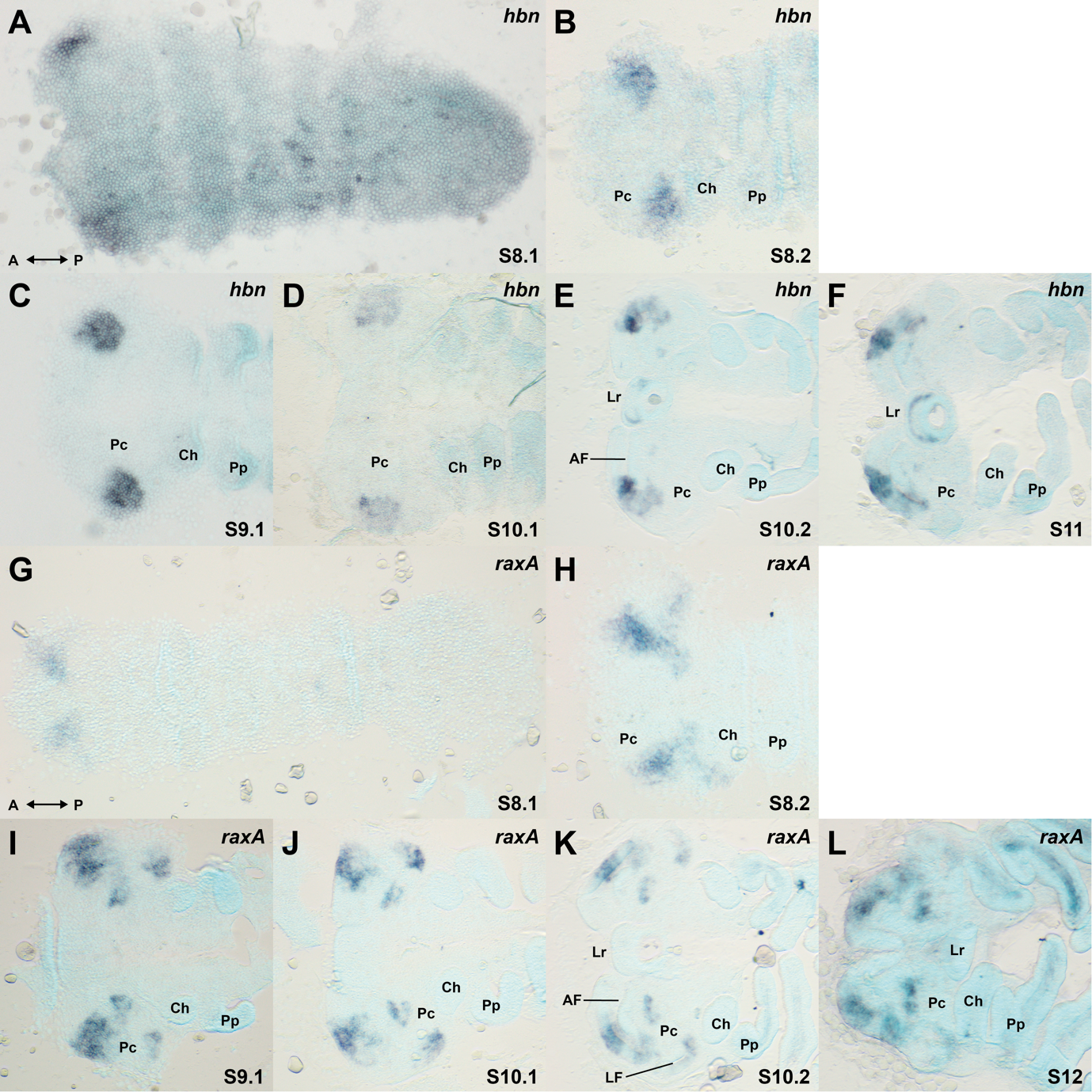

**Figure S4. Expression of *hbn* in the developing head of *P. tepidariorum* embryos.**

**A-F** Embryos stained for *hbn* transcripts at stages 8.1 **(A)**, 8.2 **(B)**, 9.1 **(C)**, 10.1 **(D)**, 10.2 **(E)**, and 11 **(F)**. **A** and **B** *hbn* was first expressed at stage 8.1, in two bilateral patches in the anterior lateral neurogenic ectoderm of the pre-cheliceral lobes. **C** and **D** From stage 10.2, *hbn* was expressed in and surrounding the lateral part of the anterior furrow (AF) and the labrum (Lr). Ch: chelicerae; Lr: labrum; Pp: pedipalps; Pc: pre-cheliceral. Embryos were flat mounted with the anterior to the left.

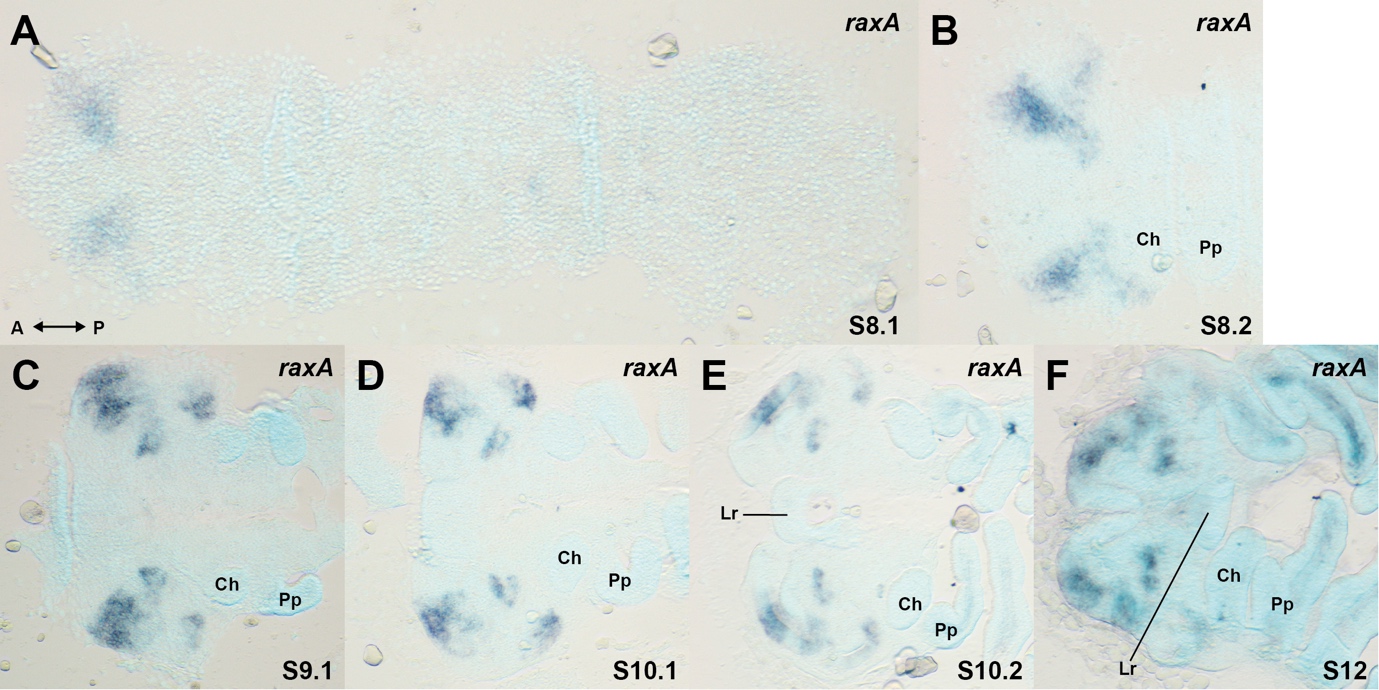

**Figure S5. Expression of *raxA* in the developing head of *P. tepidariorum* embryos.**

Embryos stained for *raxA* transcripts at stages 8.1 **(A)**, 8.2 **(B)**, 9.1 **(C)**, 10.1 **(D)**, 10.2 **(E)**, and 12 **(F)**. **A** *raxA* is first expressed at stage 8.1, in two bilateral patches in the anterior lateral neurogenic ectoderm of the pre-cheliceral lobes. **B-F** *raxA* expression subdivides into three domains, in and surrounding the lateral part of the AF, the medial pre-cheliceral lobes, and lateral of the lateral furrow (LF). Ch: chelicerae; Lr: labrum; Pp: pedipalps; Pc: pre-cheliceral. Embryos were flat mounted with the anterior to the left.

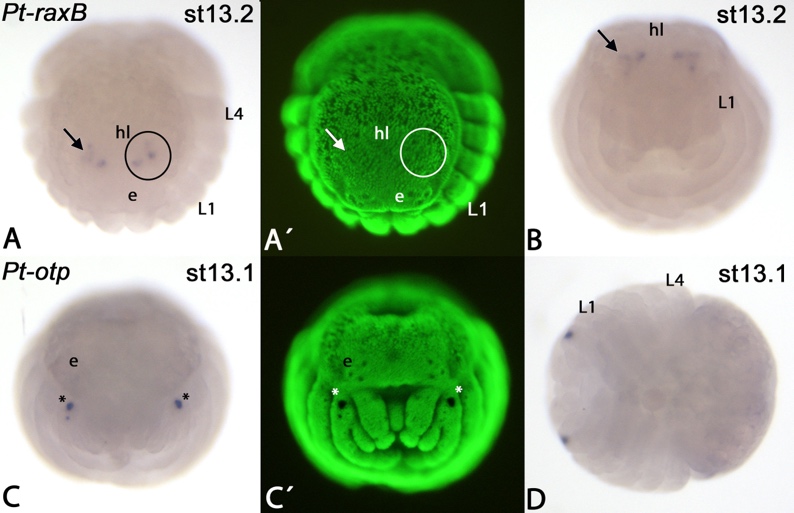

**Figure S6. Expression of *P. tepidariorum raxB* and *otp.***

The three distinct dots of *raxB* expression (arrows in **A** and **B**, encircled in **A**) do not correspond with the position of the three lateral eyes (per head hemisphere). Asterisks in panel **C** mark artificial staining of the egg teeth on pedipalps. There is no detectable mRNA expression of *otp* in all investigated developmental stages (5-13.2). Developmental stages are indicated in the upper right corner. Panels with an apostrophe represent SYBR Green staining of embryos of corresponding panels. Abbreviations: e, eyes; hl, head lobe; L, leg.

**Table S1. Primer sequences.**

| **Gene** | **Gene ID** | **Forward primer (5'‑>3')** | **Reverse primer (5'‑>3')** |
| --- | --- | --- | --- |
| *six3.2* | g25543 | **GGCCGCGG**ACAGTCCCATGTTCGTCCTT | **CCCGGGGC**GGGCTGCTACCGTGTAAATG |
| *PC1/3* | XP_015924123.1 | TCTGTTGGCAGTGCAAGTCA | **gggTAATACGACTCACTATAG**TCACCTTCCATATGACGTGGC |
| *hbn* | g8762 | **GGCCGCGG**AGGAGCGCTCTTCCTTATCC | **CCCGGGGC**GACGCTTTGCCTGAAGACAA |
| *vsx/chx* | aug3.g27186 | **GGCCGCGG**CGACCAGCAATCCAAACAGT | **CCCGGGGC**CTGACCATTGCACCGTACAG |
| *tll* | g18090 | **GGCCGCGG**CATTCTTCAACCACTGCCCC | **CCCGGGGC**TTTTCTCCAGCCTCACCAGT |
| *raxA* | g8760 | **GGCCGCGG**TGGATGTCGAAGCTCTCACT | **CCCGGGGC**TGTTGGAGGAGAGCAGGATC |
| *raxB* | g23262 | AATCGCACGATGATTGGATGC | **gggTAATACGACTCACTATAG**TCCCAGATCCTCGTTGTCCA |
| *otp* | g8767 | **GGCCGCGG**ATTCCGGCCTAGTGAAGAGC | **CCCGGGGC**TGAAAGTGACTGCGCCAAAG |
| *nkx2.1/scro* | g16180 | **GGCCGCGG**GTCCAGTTTGAGTGCTTGCA | **CCCGGGGC**GCACATAGCGGCATTGGTAA |
| *hh* | g23071 | **GGCCGCGG**GTGCCTGGCCGCATTAGTG | **CCCGGGGC**TGAGTCACCATCGAAACATC |
| *PC2* | g24083 | TCGAAATGCCACAATGCGTG | **gggTAATACGACTCACTATAG**CCTCTAGGTCCTTCACCCCA |
| *col1* | g10955 | **GGCCGCGG**AGGCAATCAGTCCTAGCGAG | **CCCGGGGC**TGATCTTGGTGCAGGTAGGG |
| *foxQ2* | g224 | **GGCCGCGG**GCGCAAATGGTAAGGGACAT | **CCCGGGGC**TTGCATGGGTGTTGAAGAGC |
| *fas2A* | g31369 | **GGCCGCGG**AAGAACTGGACGCAGGAGAA | **CCCGGGGC**CCTCACCAACAACATTGCGA |
| *Gm-fas2* | c54939_g1_i2 | GAATGCTCGGCTATGGTGGA | **gggTAATACGACTCACTATAG**TCCAGAGCCAAATTGCCGAT |
| *Gm-PC2* | c55182_g2_i11 | CGTTGAAGAAGCATGGGCAC | **gggTAATACGACTCACTATAG**CCAGCACAGGCATCAGTGTT |
